# Birth weight discordance, gene expression, and DNA methylation: A review of epigenetic twin studies

**DOI:** 10.1101/2024.11.28.625968

**Authors:** Dany Laure Wadji, Zsofia Nemoda, Chantal Martin-Soelch, Chantal Wicky

**Affiliations:** Department of Educational and Counselling Psychology, Faculty of Education, McGill University, Canada; Department of Psychology, University of Fribourg, Fribourg, Switzerland; Eating Disorders Continuum, Douglas Mental Health University Institute, Montreal, Canada; Department of Molecular Biology, Semmelweis University, Budapest, Hungary; Department of Biology, University of Fribourg, Fribourg, Switzerland

**Keywords:** Birth weight discordance, Gene expression, DNA methylation, Epigenetics, Twins studies

## Abstract

**Background:** Birth weight is considered as an important indicator of environmental conditions during prenatal development. Molecular mechanisms, including epigenetic modifications play an important role in the body’s adaptation to ever changing environmental conditions. As twin design can be used to identify the role of environmental contributions while controlling for genetic variations, numerous monozygotic twin studies have shown how adverse prenatal environment can lead to birth weight discordance (BWD).

**Objective:** An overview of the literature about epigenetic modifications associated with BWD in twins.

**Method:** We searched PubMed and Ovid MEDLINE(R) databases and included 34 papers that studied associations between BWD and DNA (hydroxy)methylation or gene expression in easily accessible samples of twin pregnancies or peripheral tissues of twins later in life.

**Results:** Researchers and clinicians still lack consensus on BWD thresholds, which vary between 15-30% depending on the type of placentation and gestational age. The gene expression twin studies measured mostly metabolism-related candidate genes in placental tissues. Only small-scale twin studies measured BWD associated gene expression patterns on genome-wide level using neonatal cells. Most DNA methylation twin studies used epigenome-level analyses, but the analysed tissue and age of sampling varied widely (blood from adults, saliva samples from children, placenta at delivery). Importantly, a handful of growth-related genes (e.g., *IGF2*, *LEPROT, ADRB3*, *GLUT3*) were associated with BWD.

**Conclusion:** Transcriptional changes of genes coding for placental glucose transporters and hypoxia-induced proteins possibly reflect compensatory processes in twin pregnancies. Epigenetic regulation of growth-related genes in the offspring offer a relevant mechanism to counterbalance adverse prenatal environment.

**Graphical abstract:** Graphical abstract/Figure 1.
(Created in https://BioRender.com) Although monozygotic twins have almost the same DNA sequence (see double stranded DNA helix in the middle), there are many molecular regulatory processes differentially affected by certain *in utero* environmental factors (such as unequal blood supply). Therefore, in a proportion of twin pregnancies, the intrauterine growth of the developing embryos is uneven, resulting in substantial birth weight difference. The underlying epigenetic alterations, such as DNA methylation changes can be long-lasting, measurable after birth and may serve as biomarkers reflecting risk for later health problems. An important technical feature is that the DNA molecule is quite stable and the methyl groups are attached covalently (shown as Me), hence methylation analyses are wide-spread in medical studies.

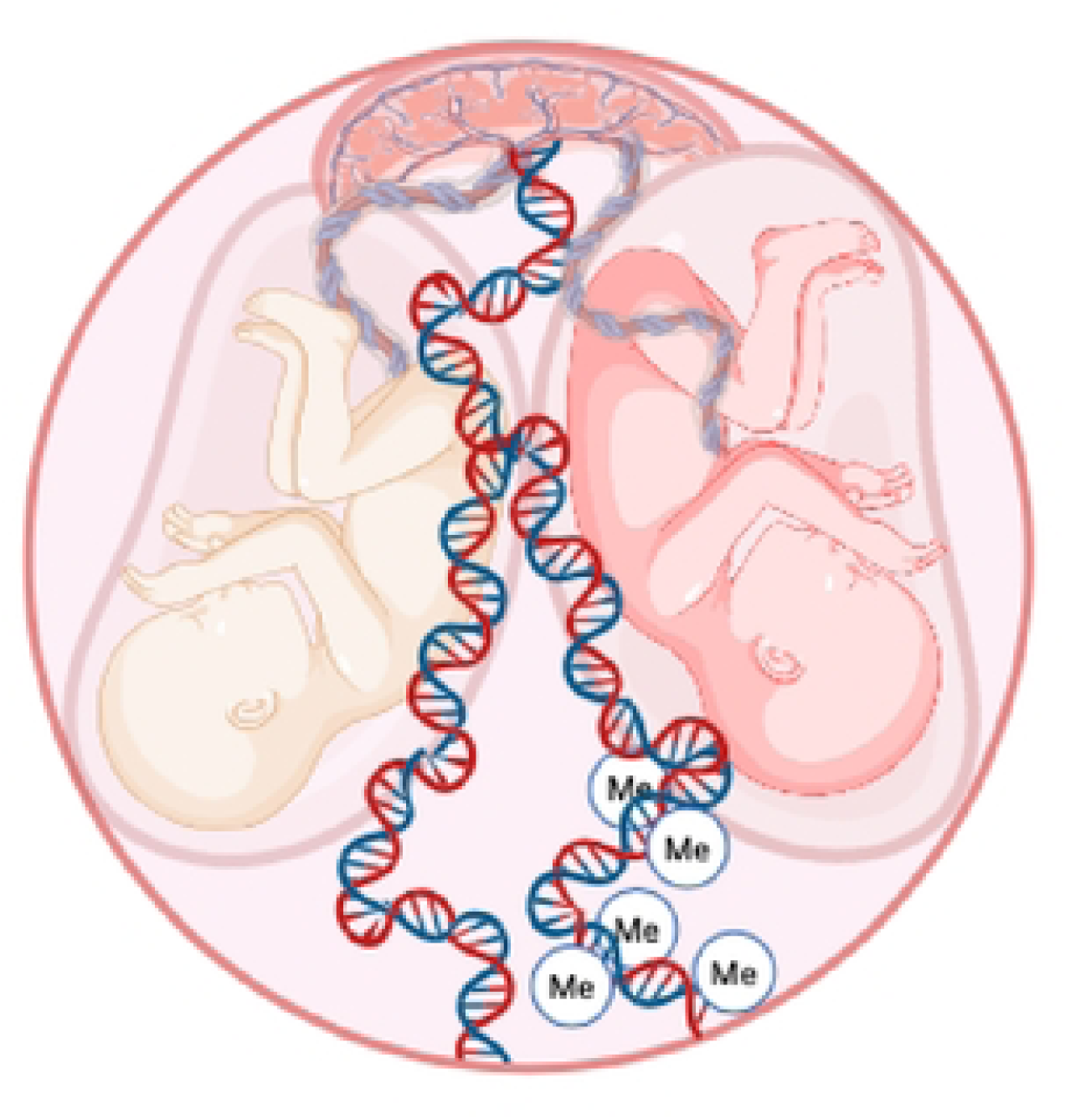

## Introduction

The proportion of multiple births (i.e., giving birth to two or more infants) has increased worldwide with an average of 16.8 per 1000 women giving birth to twins (1), possibly due to the increased use of assisted reproductive technologies, such as *in vitro* fertilization (2, 3). Multiple birth situation is likely to affect birth weight, as babies are prone to be born prematurely or before 37 weeks (4). Birth weight is considered as an important indicator of environmental conditions during prenatal development (5, 6). Birth weight discordance (BWD), i.e., substantial difference in birth weight between newborns, occurs in 10-29% of twin pregnancies (7, 8). Greater BWD has been associated with poor perinatal outcome, such as preterm birth (9), cumulative morbidity and neuro-developmental impairment in the twin with the lower weight (7, 10), intraventricular hemorrhage and cerebral palsy in the larger twin (7, 11), and intrauterine or perinatal death (12, 13). BWD is therefore of great importance for obstetrical risk assessment and for adjusting the prenatal management of such pregnancies. Numerous twin studies have assessed the underlying molecular mechanisms (e.g., gene expression or DNA methylation differences in twins), but there is no recent synthesis of this literature, which limits our ability to inform prevention and intervention strategies aimed at reducing morbidity and mortality rates in children.

### Birth weight and epigenetic markers

Molecular mechanisms, including epigenetic modifications play an important role in the body’s dynamic adaptation to environmental conditions during life, starting after conception, at implantation (14). Given that birth weight is considered as a proxy of prenatal development and is also indicative of adult health outcomes (14), epigenetic mechanisms, such as DNA methylation and histone modifications, might provide explanations on how adverse prenatal environment is associated with low birth weight (15), and how birth weight may in turn affect health later in life (16). Recent findings suggest that interindividual differences of DNA methylation patterns at certain gene regions potentially reflect distinct environmental effects (17). Therefore, epigenetic patterns associated with an indicator of growth may serve as biomarkers of adverse prenatal environment (18). Given that epigenetic modifications may be reversible, understanding these molecular mechanisms associated with birth weight could be an attractive therapeutic target that can help to minimize the long-term consequences of adverse prenatal environment.

The associations between birth weight and epigenetic markers, like DNA methylation patterns have been previously examined in several singleton studies. The most frequently measured marker in the human genome is the methylation of the cytosine base in a CpG dinucleotide (the letter “p” represents the phosphodiester bond between cytosine and guanine). Recently, patterns of additional covalent modifications, such as DNA hydroxymethylation, have been also analysed. Hydroxymethylcytosine is thought to be an important intermediate in active demethylation processes catalyzed by the ten-eleven translocation (TET) dioxygenase enzymes (19).

A large meta-analysis of epigenome-wide association studies (EWAS) conducted on 24 birth cohorts with a total number of 8,825 neonates showed that DNA methylation level at 1,029 CpG sites in 807 genes was associated with birth weight, in both positive and negative directions (45% and 55% of the sites, respectively) (18). After removing unreliably associated sites, 914 loci from 729 genes were selected for further analyses. The largest positive association was found between DNA methylation at cg06378491 in the gene body of *MAP4K2* (coding for mitogen-activated protein kinase activating kinase enzyme, which can be triggered by tumor necrosis factor alpha) – for each 10% increase in methylation at this site the birth weight was 178 grams higher. The CpG site with the largest negative association was cg10073091 in the gene body of the *DHCR24* oxidoreductase (catalyzing the reduction of the delta-24 double bond at cholesterol biosynthesis), which showed a 183 gram decrease in birth weight per 10% increase in methylation. Interestingly, 55 out of 914 differentially methylated sites were previously related to maternal smoking. An important technical point at this meta-analysis was that problematic pregnancies (e.g., mothers with preeclampsia or diabetes; multiple births, and preterm births) were excluded, hindering the applicability of the findings to any type of pregnancies.

Overall, the accumulated results from singletons’ pregnancies support the model that intrauterine environmental factors induce epigenetic alterations, which influence fetal growth and hence correlate with birth weight. These epigenetic alterations can be long lasting, but still reversible during an individual’s lifetime, therefore, the authors analysed three age groups separately. The DNA methylation data of blood samples taken in childhood (2–13 years; n = 2,756 from 10 studies) and adolescence (16–18 years; n = 2,906 from 6 studies) pointed to a few steadily affected sites, for examples in genes coding for the *GLI2* zinc finger protein and the *HOXC4* homeobox transcription factor (18). However, these birth weight associations were not detected in the adult samples (30-45 years; n = 1,616 from 3 studies).

In order to explore methylation changes during lifetime, a more appropriate type of analysis was available within the prospective Avon Longitudinal Study of Parents and Children study in the UK. Analysing 974 children with DNA methylation data from 3 timepoints (cord blood at birth, peripheral blood at age 7 and 15-17), the Accessible Resource for Integrated Epigenomic Studies identified that birth weight was associated with increased DNA methylation level at 23 CpG sites in 14 genes, but these associations could not be detected at age 7 and 17 (20). However, several associations of developmentally important genes, including the nuclear factor 1 X-type and the lymphotoxin alpha cytokine, could be found with later developmental phenotypes during adolescence, e.g., with weight, height, bone and fat mass. Later on, childhood DNA methylation levels were associated at two CpG sites with rapid weight gain, risk for obesity using independent cohort of 104 children from the Southampton Women’s Survey as a replication sample (16). Taken together, the results of singleton studies suggest that prenatal environment can affect fetal growth and birth weight through epigenetic mechanisms, and epigenetic markers may reflect risk for long-term health problems. However, the essential question remains of how to differentiate genetic effects from environmental effects in a sample of singleton pregnancies with substantial genetic heterogeneity. A twin design can be used to identify the role of unique environmental contributions (e.g., food and blood supply, unequal partitioning of nutrients) while controlling for genetic variation and shared *in utero* environment (e.g., maternal diseases, smoking or alcohol use) (21).

### The contribution of twin design

Twin design offers a unique possibility to study phenotypic variance in complex inheritance traits and diseases, because monozygotic (MZ) twins share almost identical genetic information, whereas dizygotic (DZ) twins share about 50% of genetic variants (22). Although twins share many environmental factors before birth, such as those connected to maternal habits (23), there are certain *in utero* factors, such as placental sharing and velamentous cord insertion, which create unequal food supply for the embryos, and can be related to BWD within a MZ twin pair (24). Twin studies are also valuable in testing postnatal environmental effects, if a proportion of twins are exposed to different physical, chemical or social environments. Therefore, twin design can be used in various models, e.g., the Developmental Origins of Health and Disease theory hypothesizing that exposure to prenatal and early life events during critical periods of development and growth may shape adult health and vulnerability to disease later in life (25). It is argued that epigenetic processes can account for these long-lasting effects. Consequently, many studies examined associations between gene expression or DNA methylation patterns and BWD using twin models. Here we summarize the state of research about epigenetic changes associated with BWD in twins. Although BWD can be used as a continuous variable (scale) within twin pair comparison in the association analyses, many researchers compared low vs. high BWD pairs, but the grouping methods differed in the epigenetic studies. Therefore, we will first discuss the epidemiological twin studies creating threshold values of BWD, then we present the state of research on BWD, gene expression and DNA (hydroxy)methylation.

### Factors associated with birth weight discordance and threshold values

Substantial difference in birth weight of twins is argued to be related to prenatal factors acting asymmetrically on the two embryos, affecting the intrauterine growth of one of them while the growth of the larger twin is apparently not affected (26). Depending on the type of placentation, i.e., monochorionic (MC) or dichorionic (DC) pregnancies, several intrinsic and extrinsic factors have been associated with BWD (13). Intrinsic factors can be attributable to the twin pregnancy situation or to the twins themselves. For example, unequal placental sharing (27), velamentous or other abnormal cord insertion (28), and twin-twin transfusion syndrome (TTTS), i.e., when blood moves from one twin to the other, can lead to fetal growth restriction (FGR) in MC twin pregnancies (9, 26). The TTTS happens approximately 15% of MC twin pregnancies, when the placenta cannot provide enough oxygen and nutrients for both fetuses, and subsequently blood flows from one of the twins to the other (creating donor and recipient twin). The ultrasound signs therefore polyhydramnios in the recipient twin (due to increased blood flow, and subsequent urination into the amniotic fluid) and oligohydramnios in the donor twin, who grows slower, because of less blood flow (29). Umbilical blood vessel abnormalities can cause growth retardation even is DC pregnancies (see for examples, 30; 31). Hormone imbalances, such as lower adiponectin concentration can also result in growth restriction in discordant twins (32). Extrinsic factors include maternal medical conditions, such as high blood pressure, preeclampsia, hepatitis C infection, and delayed childbearing (13, 33, 34), socioeconomical status of the family, reflected by parental education (35), or maternal habits, e.g., smoking during pregnancy (33).

The discordance of birth weights (in %) is calculated as = (weight of larger twin - weight of smaller twin) / weight of larger twin × 100. The BWD thresholds are used for the prediction of perinatal mortality and morbidity. However, there is still no consensus among researchers and clinicians on BWD thresholds which have been ranging between 15-40% depending on type of twins, type of placentation (chorionicity), and sex of twins (see Table 1). Important technical differences come from the study design: population-based studies with more than thousands of twin pairs used only birth and death certificates and did not have data on zygosity and chorionicity, therefore, specific subsets were analyzed separately (see for example 36). Also, the threshold values of gestational age ranged widely, most studies included pregnancies after 24 weeks of gestation, but a few selected only term pregnancies (e.g., 37), or excluded fetal death cases (9). Therefore, the inconsistent results about threshold of BWD may be due to the fact that most research groups have relied basically on the level at which adverse perinatal and postnatal outcomes are more likely to occur in their specific subset of twin pregnancies.

**Table 1:** Threshold values for birth weight discordance used in epidemiological twin studies. At the sample description the number of monozygotic (MZ) or dizygotic (DZ) twin pairs are shown. Wherever possible, the number of monochorionic (MC) and dichorionic (DC) pregnancies are indicated. Only percentages are indicated when chorionicity data was partially available. Most studies excluded monoamniotic twins, major congenital abnormalities (except for persistent ductus arteriosus in case of preterm birth), and twin-twin transfusion syndrome (TTTS); * shows where information on possible TTTS could not be obtained. Fetal death was defined as intrauterine death of the fetus ≥ 20 weeks of gestation, neonatal death was defined as death of the infant within 28 days after birth, whereas combined perinatal mortality included loss of twin(s) after 24 completed weeks or newborn weighting at least 500 g. Hospital death means death of newborn before discharge or within 120 days for infants still hospitalized. Preterm birth was defined as a birth that takes place before 37 weeks of pregnancy have been completed, if not otherwise specified by the study protocol (see details in the last column). *Other abbreviations:* BW: birth weight; BWD: birth weight discordance; CI: confidence interval; GA: gestational age; NICU: neonatal intensive care unit; NRDS: neonatal respiratory distress syndrome; OR: odds ratio; RR: relative risk; SGA: small for gestational age (less than the 10^th^ percentile); SES: socioeconomic status; twin A: the first baby delivered at birth

| Authors, birth cohort | Sample characteristics, exclusion criteria | Covariates (infant, maternal and pregnancy related variables) | BWD cutoff | Outcome variables associated with BWD |
| --- | --- | --- | --- | --- |
| Ye et al. (2022)<br>Foshan, China,<br>2012-2018 | 1,946 DC and 402 MC twin pairs without TTTS (total 2,348 pairs), GA $\geq 26$ weeks | GA, SGA, sex, BW; maternal age, ethnicity, marital status; chorionicity, use of assisted reproductive technology, parity, obstetric complications, mode of delivery | 15-30% depending on the variable | BWD $\geq 15\%$ was associated with NICU admission (OR = 1.54; 95% CI: 1.23 - 1.93), a BWD $\geq 20\%$ for NRDS (OR = 2.20; 95% CI: 1.38 - 3.51) and a BWD $\geq 30\%$ for ventilator support (OR = 2.39; 95% CI: 1.25 - 4.57). |
| Chen et al. (2020)<br>US, 1995-2000 | 98,588 sex-discordant (hence DZ) twin pairs | GA; maternal age, race, marital status, education, parity, smoking during pregnancy, pregnancy-associated hypertension | 40% | BWD $\geq 40\%$ was associated with composite adverse neonatal outcome combining NRDS, meconium aspiration syndrome, asphyxia at delivery, neonatal seizures (OR = 1.6; 95% CI: 1.1 - 2.3) and fetal death at GA 33-34 gestational weeks (OR = 17.3; 95% CI: 9.2 - 32.7). |
| Boghossian et al. (2019)<br>US, 1994-2011 | 8,114 preterm twins with very low birth weight (401-1500 g) or gestational age 22–28 | GA, SGA, sex; birth year, study center; maternal ethnicity/race, antenatal steroid use | 30% (separately for smaller | In smaller twins: BWD $> 30\%$ was associated with increased risk for hospital death (OR = 3.2; 95% CI: 2.1-4.8), retinopathy (OR = 2.4; 95% CI: 1.1 - 5.4), bronchopulmonary dysplasia (OR = 2.3; |
|  | weeks, TTTS twins were excluded if it was the cause of death |  | and larger twins) | 95% CI: 1.6 - 3.6), necrotizing enterocolitis (OR = 2.1; 95% CI: 1.3 - 3.4).<br>In larger twins: BWD > 30% was associated with risk for patent ductus arteriosus (OR = 1.5; 95% CI: 1.1 - 1.9) and OR ~ 5 for later neurodevelopmental impairment, blindness and cerebral palsy (assessed at age 1 year). |
| Kim et al. (2019) US, 2012-2014 | 27,276 twin pairs, if twin A was delivered vaginally in cephalic presentation at GA between 36-40 weeks | sex; maternal age, race/ethnicity, marital status, SES, obesity; parity, breech delivery of the second twin | 20% | BWD > 20% was associated with composite adverse neonatal outcome (e.g., 5-minute Apgar < 7, NICU admission, mechanical ventilation > 6 hours, neonatal seizure) (adjusted OR: 1.46; 95% CI: 1.29 - 1.65). |
| Vedel et al. (2017) Denmark, 2004-2008 | 2,378 DC and 355 MC twin pairs without TTTS (total N = 2,733 pairs), GA ≥ 24 weeks | GA, SGA, sex | 15% | BWD > 15% was associated with induced preterm delivery at GA < 34 weeks (OR = 1.71; 95% CI: 1.11 - 2.65); increased risk of infant mortality after 28 days (OR = 4.69; 95% CI: 1.07 - 20.45); risk for admission to NICU for more than a week (OR = 1.53; 95% CI: 1.21 - 1.95). |
| Breathnach et al. (2011) Ireland, 2007-2009 | 789 DC and 188 MC (14 with TTTS, total N = 963 pairs without TTTS), GA ≥ 24 weeks | GA, SGA, sex, chorionicity, birth order, | 18% | BWD > 18% was associated with adverse perinatal outcome in MC twin pairs without TTTS (hazard ratio = 2.6; 95% CI: 1.6 - 4.3) and in DC twin pairs (hazard ratio = 2.2; 95% CI: 1.6 - 2.9). |
| Amaru et al. (2004)* New York, US, 1992-2001 | 1,318 twin pairs (68% had chorionicity data, TTTS is not mentioned), GA ≥ 24 weeks | GA, chorionicity, oligohydramnios (sign of TTTS), antenatal steroid use, preeclampsia | 20% | BWD > 20% was associated with cesarean delivery (OR = 1.87; 95% CI: 1.22 - 2.87), adverse neonatal outcomes, like very low birthweight (OR = 8.73; 95% CI: 3.15 - 24.19), NICU admission (OR = 3.26; 95% CI: 1.97 - 5.40), neonatal oxygen requirement (OR = 1.71; 95% CI: 1.00 - 2.90), hyperbilirubinemia (OR = 1.69; 95% CI: 1.07 - 2.66). |
| Ananth et al. (2003) & Demissie et al. (2002)* US, 1995-1997 | 269,287 twin pairs were divided to same sex and different sex (hence DZ) groups due to the lack of zygosity and chorionicity data | GA, SGA, sex; maternal age, education, illnesses (e.g., chronic hypertension, diabetes), parity, prenatal care utilization; placenta previa | 20-40% depending on sex and size of the twins | Increasing BWD was associated with overall neonatal deaths (OR = 1.29; 95% CI: 0.99 - 1.68) among larger but not smaller twin of the same sex (this association was not found among twins of opposite sex).<br>BWD ≥ 20% in same sex twin pairs (RR = 1.2; 95% CI: 1.1 - 1.4) and BWD ≥ 40% in different sex twin pairs (RR = 2.2; 95% CI: 1.7 - 2.8) was associated with risk for placental abruption. |
| Hartley et al. (2002)*<br>Washington, US, 1987-1999 | 7,803 twin pairs, chorionicity and TTTS information was not available | GA, SGA | 25% | BWD > 25% was associated with perinatal death (RR = 2.2; 95% CI: 1.5 - 3.1). |
| Redman et al. (2002) Michigan, US, 1995-2000 | 173 twin pairs (14% MC), GA > 23 weeks | GA, chorionicity | 30% | BWD > 30% was associated with cesarean section for non-reassuring fetal status, NICU admission, pH < 7.1 in umbilical artery, and 5-minute Apgar score < 7 ($p < .0001$). |
| Victoria et al. (2001) Missouri, US, 1993-1995 | 288 DC and 89 MC twins (5 with TTTS), GA > 24 weeks (N = 377 pairs) | GA, SGA, chorionicity maternal age, poor obstetric history | 25% | BWD > 25% was associated with increased odds for adverse neonatal outcomes like preterm birth < 36 weeks GA (OR = 5.05; 95% CI: 1.05 - 27.22), NICU stay of more than 10 days (OR = 3.28; 95% CI: 1.17 - 9.30) in severely discordant DC and MC twin pairs compared to severely discordant MC twin pairs. |
| Cooperstock et al. (2000) Missouri, US, 1978-1990 | 9,931 twin pairs, excluding pregnancies with fetal death (in either one twin) | GA, SGA, sex; maternal age (being a teenage mother), race, SES, parity, cigarette smoking | 30% | BWD > 30% was associated with preterm twin births at GA < 32 weeks (OR = 1.77; $p < 0.03$). |

The largest epidemiological twin studies (using nation-wide datasets) divided their analyses into same sex and different sex groups, because zygosity and chorionicity data was not available. They found BWD cut-off values of 15% for same sex twins and 30% for different sex (i.e., DZ and hence DC) twins conferring the greatest risks for fetal and neonatal death, and preterm birth (36). Demissie et al. (38) showed that fetal mortality rates differed between the smaller and larger twins of the opposite sex (adjusted odds ratio > 2.7 from BWD > 20% among the smaller twins, but no consistent increase in the odds among the larger twins), whereas same sex twins had similar odds for fetal death (adjusted odds ratio > 1.7 from BWD > 20%). Later, Y. Chen et al. (39) included only mixed gender twins as a surrogate data for chorionicity. Among sex-discordant premature DZ twins a cut-off value of 40% was associated with adverse neonatal outcomes (for details, see Table 1).

The earliest clinical studies included MC twins with TTTS, when blood flows from one of the twins to the other (creating donor and recipient twin). In this setup, Victoria et al. (40) found that severe BWD (> 25%) was more frequent in MC than DC twins, explained mostly by small placenta and umbilical cord abnormalities. Later, Breathnach et al. (11) showed that difference in BWD cut-off values associated with perinatal mortality and morbidity among DC and MC twins (18% *vs* 15%) disappeared when MC twins with TTTS were excluded from the analyses. Therefore, most research groups examined twin pairs without TTTS and also excluded the rare monoamniotic twin pregnancies in order to rule out exceptional cases leading to extreme adverse neonatal outcomes. It has to be noted, that severe TTTS cases can be treated with laser surgery during pregnancy, which can normalize the conditions for both fetuses (41).

One of the first studies was conducted in the United States by Redman et al. (42) who reported that 95th percentile (BWD > 30%) was the threshold most predictive of adverse neonatal outcomes, like cesarean section, umbilical artery and neonatal intensive care unit admission. Later, a prospective study conducted in Ireland by Breathnach et al. (2011) showed lower threshold; adverse neonatal outcome was increased with BWD exceeding 18%. In order to rule out other confounding factors, such as separate vs common placenta between DC and MC twins, Amaru et al. (43) controlled for chorionicity in their analyses, and reported that 20% BWD cut-off value was associated with cesarean delivery and adverse neonatal outcomes, like neonatal intensive care unit admission, oxygen requirement and hyperbilirubinemia. A Danish study found a threshold value of 15% in DC twins where low birth weight was associated with mortality (44). Recently, in a Chinese cohort Ye et al. (45) reported cut-off values between 15-30% associated with different adverse neonatal outcomes.

In twin cohorts of unknown chorionicity and much larger sample size, BWD cut-off values ranged from 20 to 40%. For example, threshold values between 30-40% were reported by an early study by Cooperstock et al. (9) who used a large US database from 1978 until 1990 to study the association between BWD and preterm birth. Using a population-based retrospective analysis of birth certificates and fetal/infant death certificates in the state of Washington, Hartley et al. (46) found BWD cut-off value of 25% indicative for perinatal and neonatal mortality. Boghossian et al. (7) examined BWD in premature, very low birth weight twins and found that most adverse outcomes including death before discharge from the hospital, necrotizing enterocolitis, severe retinopathy, and bronchopulmonary dysplasia, were more common among the smaller twins with the highest BWD (> 30%). The larger twins with the highest discordance level had increased odds of patent ductus arteriosus, moderate-to-severe cerebral palsy, and blindness. Even among term babies born between 36-40 weeks, Kim et al. (37) reported 20% BWD cut-off value associated with worse perinatal outcomes in a cohort of twins with vertex twin A delivered vaginally. Taken together, BWD cut-off values indicative for adverse perinatal outcomes are around 20-30%, their determination depends on the chorionicity, the sex of twins, the gestational age, and the presence of TTTS.

## Materials and Methods

We searched for papers studying the association between BWD and gene expression or DNA methylation using two databases up to the date April 19, 2024. The following keywords were used in the PubMed database: “birth weight discordance” / “discordant growth” / “twin birth weight difference” & “gene expression” / “DNA methylation” / “DNA hydroxymethylation”. At the Ovid MEDLINE(R) we used “birth weight” / “growth restriction” & “twin” & “gene expression” / “DNA methylation” / “DNA hydroxymethylation” keyword combinations in order to include wider range of reports. The automatic search was performed independently by two authors (DW and ZN), who reviewed all abstracts independently. The inclusion criteria for our review were the followings: (i) published in peer-reviewed journal, (ii) examine BWD in human twins, (iii) focus on epigenetic mechanisms, like DNA methylation or gene expression. Disagreements about exclusion were resolved by discussion. Abstracts and full text of the potential papers were reviewed.

The PubMed search resulted in 36 papers with human data, the Ovid MEDLINE(R) search resulted additional 6 papers on human samples (another 2 studies were conducted on animals). Finally, the automatic search was completed with a manual search from the reference lists of previous systematic reviews, meta-analyses, and the retrieved articles (adding one more paper). The combined search yielded 43 papers (after removing duplicates), from which 9 were excluded, because the report used the same twin cohort as in Gao et al. (47) paper, or it did not include original research results (2 reviews), or there was no BWD analyses conducted with gene expression or DNA (hydroxy)methylation (6 papers). At the end, 34 independent papers were eligible for our review: 19 had gene expression, 21 had DNA methylation, and 2 had DNA hydroxymethylation measurements (8 of them were overlapping having both gene expression and (hydroxy)methylation data).

## Results

### Gene expression and birth weight discordance

The majority of gene expression studies selected *a priori* candidate genes implicated in metabolism or cardiovascular development or fetal growth (summarized in Table 2). For example, Gao et al. (47) used placental samples to examine the expression of the human endogenous retrovirus group W member 1, envelope gene (*HERVWE1*) known to be highly expressed in syncytiotrophoblasts when creating the syncytial layer of the placenta. In addition, HERVWE1 is involved in several endocrine hormone pathways, such as the human chorionic gonadotrophin and somatomammotropin pathways, that promote fetal growth. In addition to 56 MZ DC twin pairs, they collected data from 10 singleton pregnancies (for comparisons’ reason, none of them had growth restriction, and all of them were delivered by cesarean section between 30-36 gestational weeks). Significant differences were observed only among the 21 discordant twin pairs with at least 20% BWD: *HERVWE1* mRNA and protein levels were increased in the smaller twins compared to the larger twins, with an accompanying decrease in DNA methylation level at the gene’s promoter region. Furthermore, the *HERVWE1* transcript level was negatively correlated with birth weight in all the groups, which was attributed to the unique characteristics of the study population, consisting of discordant (but not FGR) twins with birth weights within the normal range.

**Table 2:**
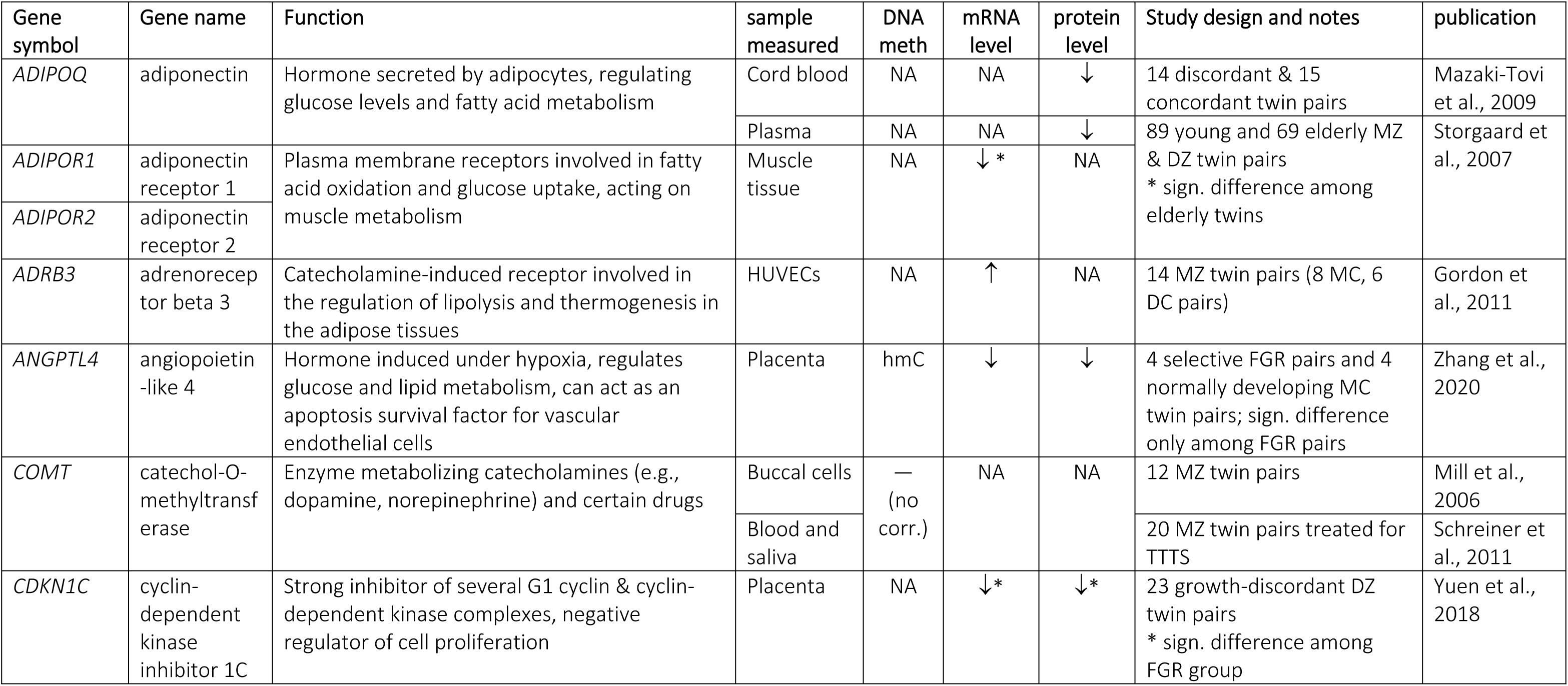
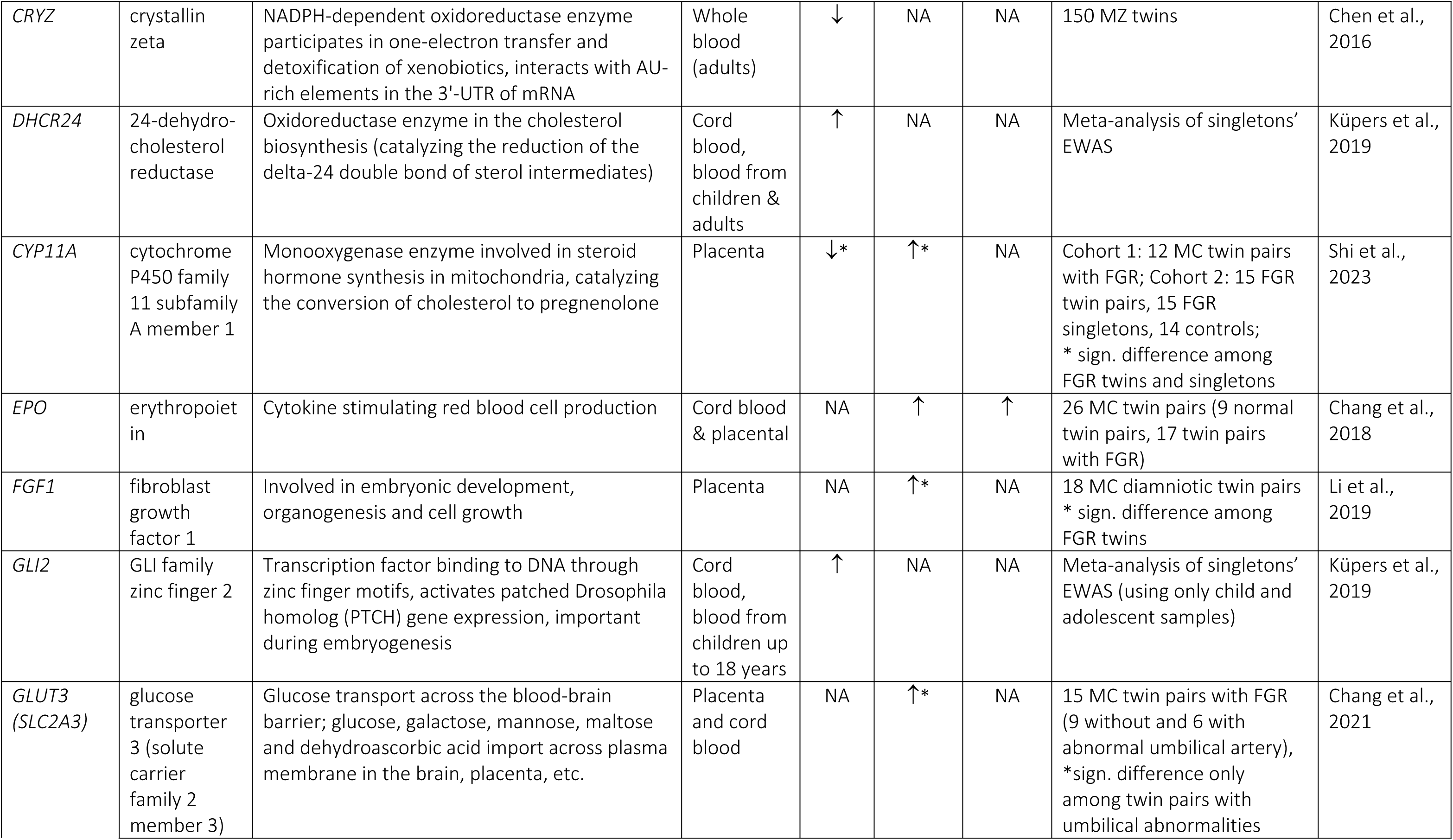
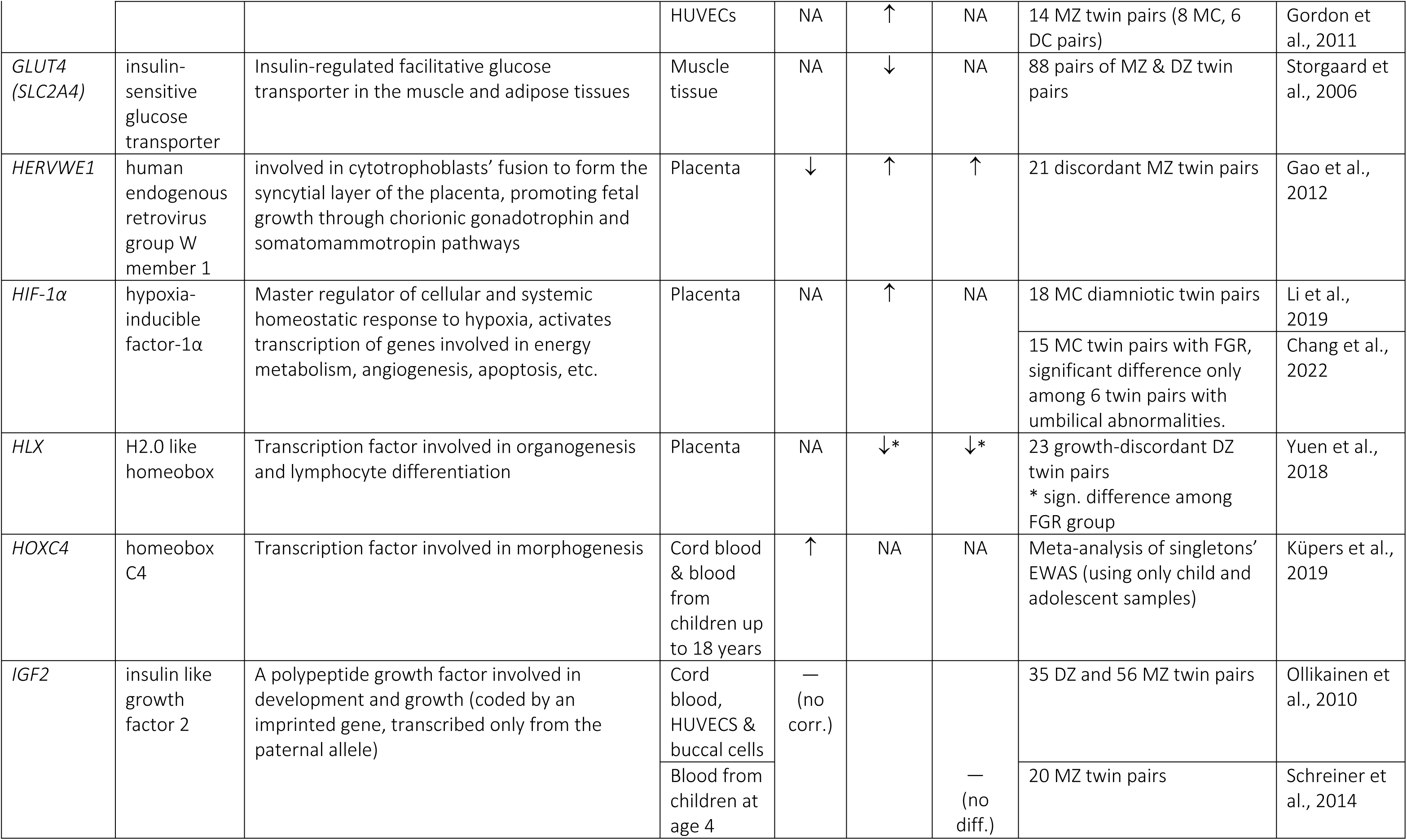
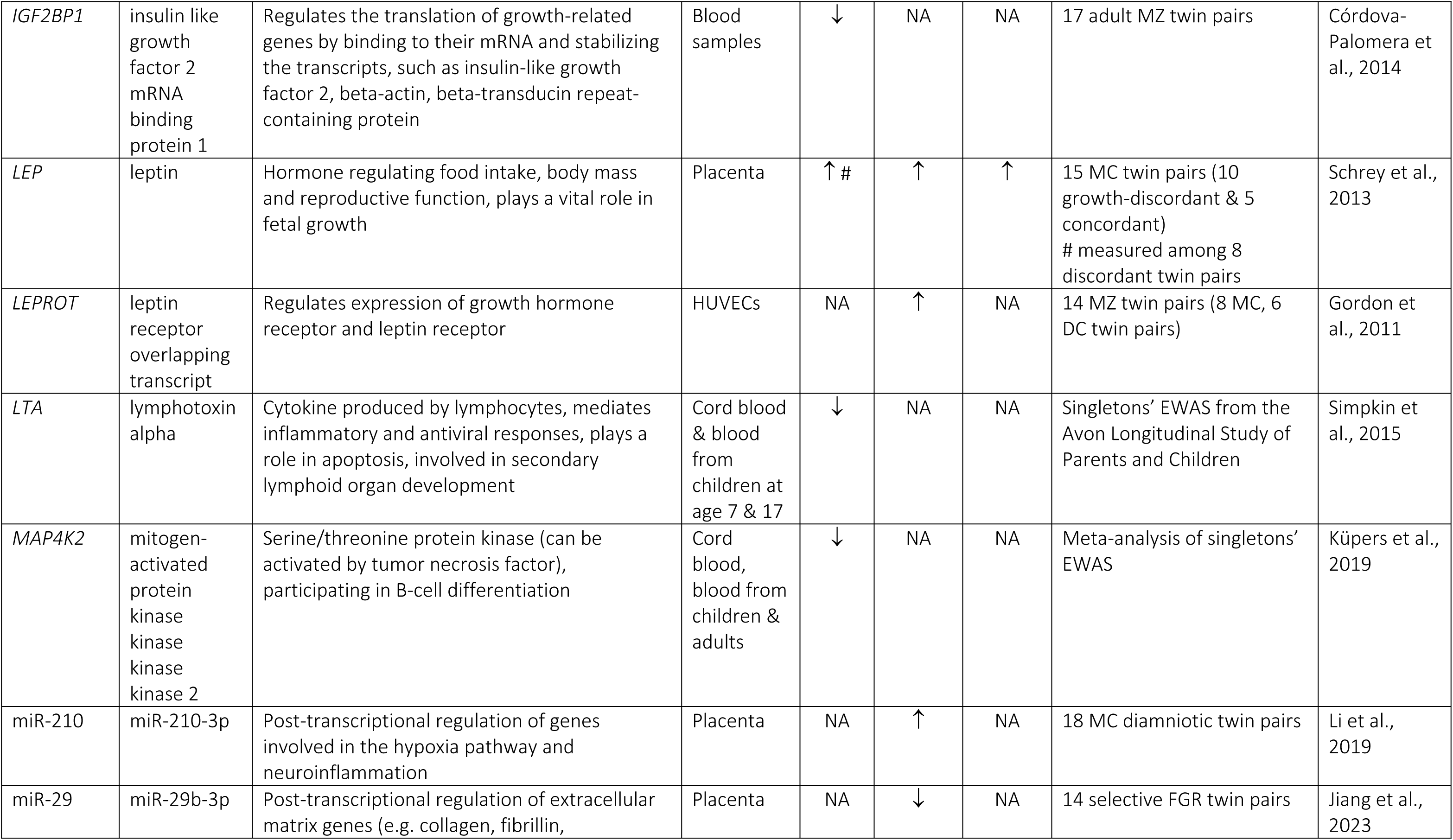
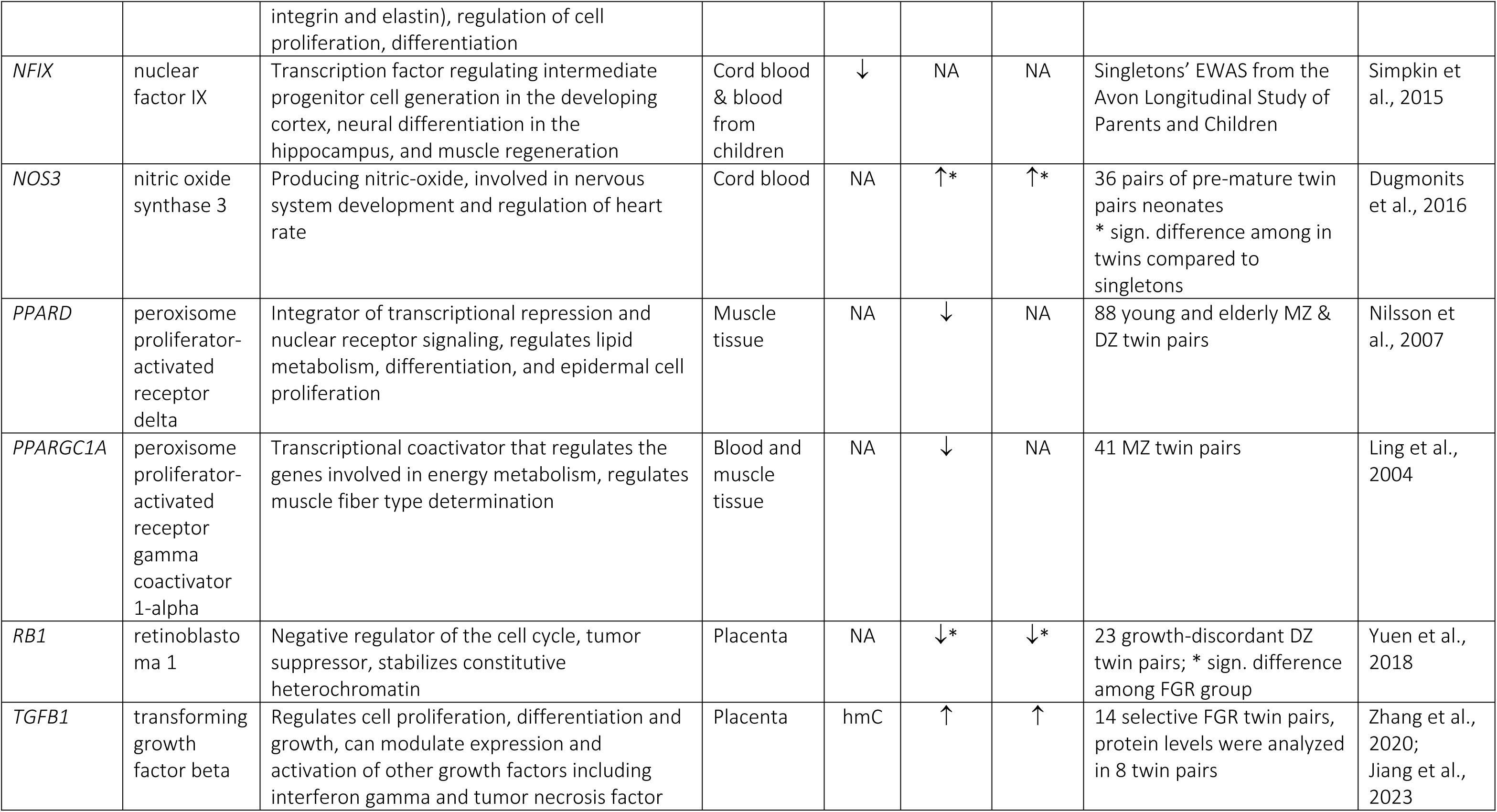
Epigenetic changes associated with birth weight discordance, resulting in lower birth weight. Gene expression and DNA (hydroxy)methylation differences are shown by arrows according to changes in the smaller twins compared to the larger twins. The associations between birth weight and DNA methylation, mRNA and protein levels observed in singleton birth cohorts are given to reflect changes resulting in lower birth weight. *Abbreviations:* DC: dichorionic; DZ: dizygotic; EWAS: epigenome-wide association studies; FGR: fetal growth restriction; hmC: hydroxymethylcytosine; HUVECs: human umbilical vein endothelial cells; MC: monochorionic; MZ: monozygotic; NA: not assessed; no corr.: no correlation; sign.: significant

Some researchers examined twin pregnancies with selective FGR, i.e., when the smaller twin’s birth weight is less than 10^th^ percentile in the third trimester, with inter-twin BWD > 20-25%. For example, Schrey et al. (48) investigated 88 angiogenesis-related genes in placental samples of 10 severely growth-discordant MC twins (BWD ≥ 20%, without TTTS) and 5 growth-concordant MC twins. They found 3 genes showing mRNA expression differences between the smaller and larger twins, and confirmed at the protein level that leptin expression, which regulates food intake and plays a vital role in foetal growth, was increased in the smaller twins of the discordant growth group. The DNA methylation analysis of the leptin gene promoter showed higher methylation in the smaller twins, potentially resulting in a reduced binding of a repressor type of transcription factor at this regulatory region.

Using a similar study design, Chang et al. (49) measured cord blood erythropoietin level and placental erythropoietin gene expression in MC twins delivered through elective cesarean section (9 normal pairs, 9 pairs with selective FGR but without umbilical artery abnormalities, 8 pairs with FGR and with artery abnormalities), and calculated ratio of erythropoietin level of the smaller / larger twin within each twin pair. They found good correlation between fetal plasma erythropoietin and placental expression ratios in the whole sample. Twins with FGR and umbilical artery abnormalities had the highest plasma erythropoietin level (compared to the other two groups), possibly compensating for the suboptimal circulation, increasing erythropoiesis to enhance the oxygen-carrying capacity of their blood. The smaller twins of pregnancies with umbilical artery abnormalities had two times higher cord plasma erythropoietin level and placental expression compared to their co-twins, who had significantly elevated erythropoietin level compared to the other groups (they also had significantly shorter gestation period and smaller birth weight compared to the other twins). The same group later examined placental glucose transporters’ expression. Only twins with FGR and umbilical artery abnormalities differed from the others at the glucose transporter 3 (*GLUT3/SLC2A3*) level (50). Using placental mesenchymal stem cells derived from MC pregnancies with selective FGR, Chang et al. (51) finally showed that the group of growth-restricted fetuses had lower capacity to increase the placental glucose transporter expression under hypoxic conditions compared to their appropriate-for-gestational-age co-twins.

An Australian study investigated expression level of *HLX* homeobox gene coding for a transcription factor and its downstream target genes (e.g., retinoblastoma and cyclin dependent kinase inhibitors) in placental samples of 23 DZ twins (52). Decreased expression of *HLX* and cyclin dependent kinase inhibitor 1C (*CDKN1C*) was detected in the FGR group at both the mRNA and protein levels, relative to the normal group, potentially reflecting slower development and subsequently retarded fetal growth. Another type of regulation was studied by Li et al. (53), analysing microRNAs in 18 placenta sections of diamniotic MC twins (excluding TTTS, chromosomal abnormalities, and maternal diseases). Increased expression of miR-210-3p was found in the FGR group, which can lead to decreased expression of fibroblast growth factor 1. Their *in vitro* experiments supported the following model: under hypoxic conditions, level of hypoxia-inducible factor-1α (*HIF-1α*) is increased, causing an increased miR-210-3p expression, which in turns inhibits fibroblast growth factor 1 gene expression. Altogether, this leads to impaired proliferation of the trophoblast cell line.

*In vivo* assays were also used in twin studies to decipher birth weight-related epigenetic mechanisms leading to health problems later in life, since metabolic diseases, such as diabetes mellitus are often linked to prenatal risk factors (54). In a series of papers, almost a hundred healthy, full-term twin pairs (born 40 ± 3 gestational weeks) from the Danish Twin Register were analysed for metabolic markers in muscle biopsy samples taken before and during insulin infusion (testing their insulin sensitivity). Among the 41 MZ twin pairs with gene expression data, birth weight was positively associated with the insulin-stimulated expression of *PPARGC1A* (peroxisome proliferator-activated receptor gamma coactivator 1-alpha) and glucose oxidation rate (55). Significant positive association was found between birth weight and both baseline and insulin-stimulated glucose transporter 4 level measured in the muscle (56). Later, they showed similar positive association of birth weight and peroxisome proliferator-activated receptor delta muscle expression (57). In addition, plasma samples were analysed for adiponectin, which is an insulin-sensitizing adipokine acting on muscle metabolism (58). Nongenetic influence of birth weight was observed on both plasma adiponectin level and muscle expression of adiponectin receptor 1 among MZ twins. Interestingly, birth weight was also associated with *in vivo* measures of glucose metabolism in these twins, highlighting the fetal programming effects on glucose homeostasis, and their possible roles in the pathogenesis of type 2 diabetes later in adulthood (59) (for more details see Table 2).

Analysing the role of oxidative stress in delayed fetal development and subsequently low birth weight, Dugmonits et al. (60) measured the levels of reactive oxygen species in red blood cells from twin and singleton neonates. Higher levels of hydrogen peroxide and nitrate, and consequently elevated protein and lipid damages were reported in twin neonates compared to singletons. Expression levels of nitric oxide synthase and other enzymes of the antioxidant defence system (including catalase, superoxide dismutases and haemoxygenases) showed that premature twins with lower birth weights had the lowest antioxidant capacity.

One of the first genome-wide studies analysing associations between BWD and gene expression without *a priori* hypotheses used multiple tissue types from the Peri/postnatal Epigenetic Twins Study. Gordon et al. (61) showed significant negative correlation between low birth weight and expression of genes related to metabolism and cardiovascular function. They found 41 genes significantly linked to birth weight in human umbilical vein endothelial cells (HUVECs) analysed from 14 twin pairs. Comparing cord blood mononuclear cells, only DC twins showed differential expression at 342 genes, highlighting the possibility that MC twins, who share the same placenta, have lower gene expression difference. They highlighted three genes involved in fetoplacental growth and metabolism, which expression in HUVECs was negatively correlated with birth weight: adrenoreceptor beta 3, glucose transporter 3, and the leptin receptor overlapping transcript, which has an important regulatory role in the cell surface expression of growth hormone and leptin receptors (for more details, see Table 2). These results suggest compensatory mechanisms at the expression of genes involved in fetoplacental growth and supplying energy.

Recently, a Chinese group compared placental mRNA levels and DNA hydroxymethylation patterns of 4 twin pairs with selective FGR to 4 normally developing MC twin pairs. Their mRNA-sequencing data showed 81 up-regulated and 614 down-regulated genes in the placental shares of the FGR group compared to the co-twins. Analysis of the overlapping genes with differential mRNA expression and DNA hydroxymethylation levels pointed to the involvement of angiopoietin-like 4 (*ANGPTL4*) gene. Based on functional assays, the authors suggested a model in which hypoxia through the HIF-1α pathway changed the *ANGPTL4* gene promoter methylation pattern, leading to decreased *ANGPTL4* level (62). Further exploring the molecular process of this regulatory pathway, Jiang et al. (63) measured the TET enzyme family (proposed to be the main catalysators of the active DNA demethylation processes), as well as their activators, inhibitors, and downstream targets in placental samples of 14 pregnancies with selective FGR (without TTTS or other reversed arterial perfusion). They identified a downregulated microRNA (miR-29b-3p) causing increased TET2 enzyme activity and subsequently increased transforming growth factor beta 1 (TGFB1) level in the FGR samples. Since TGFB1 is an important regulator of cell proliferation and differentiation, the authors highlighted the role of miR-29b-3p-TET2-TGFB1-smad axis leading to reduced trophoblast invasion and to increased apoptosis in growth retardation.

Altogether, using neonatal cells from cord blood or placental tissues at birth, gene expression studies provided evidence that prenatal environment can affect expression of genes involved in fetal growth (e.g., *LEPROT, HERVWE1, TGFB1*), oxygen and glucose delivery (e.g., erythropoietin, glucose transporters), and hypoxia-signaling cascade (e.g., HIF-1α). However, it has to be noted that most reports did not have replication cohort and used relatively small sample size.

### DNA methylation and birth weight discordance

Except for a few studies analysing placental tissues, most research groups investigated BWD associated DNA methylation patterns in cord blood or later in life using peripheral blood or saliva samples. The most recent analyses used sequencing or the Illumina EPIC array with over 850 thousands of CpG sites, whereas the majority of published data is based on the Illumina 450k array (see details in Table 3). Researchers also started to examine another covalent modification of the genomic DNA, the hydroxymethylation, which can serve as a marker for active demethylation, hence gene activating processes. The above-mentioned Chinese workgroup analysed hydroxymethylated regions of the genome in placental samples of pregnancies with selective FGR and compared them to expression changes. In this way they highlighted the aberrant hydroxymethylation pattern at the *ANGPTL4* gene, resulting in reduced protein expression in the smaller placental shares, leading to reduced trophoblast functions, and subsequent suboptimal growth (62).

**Table 3:**
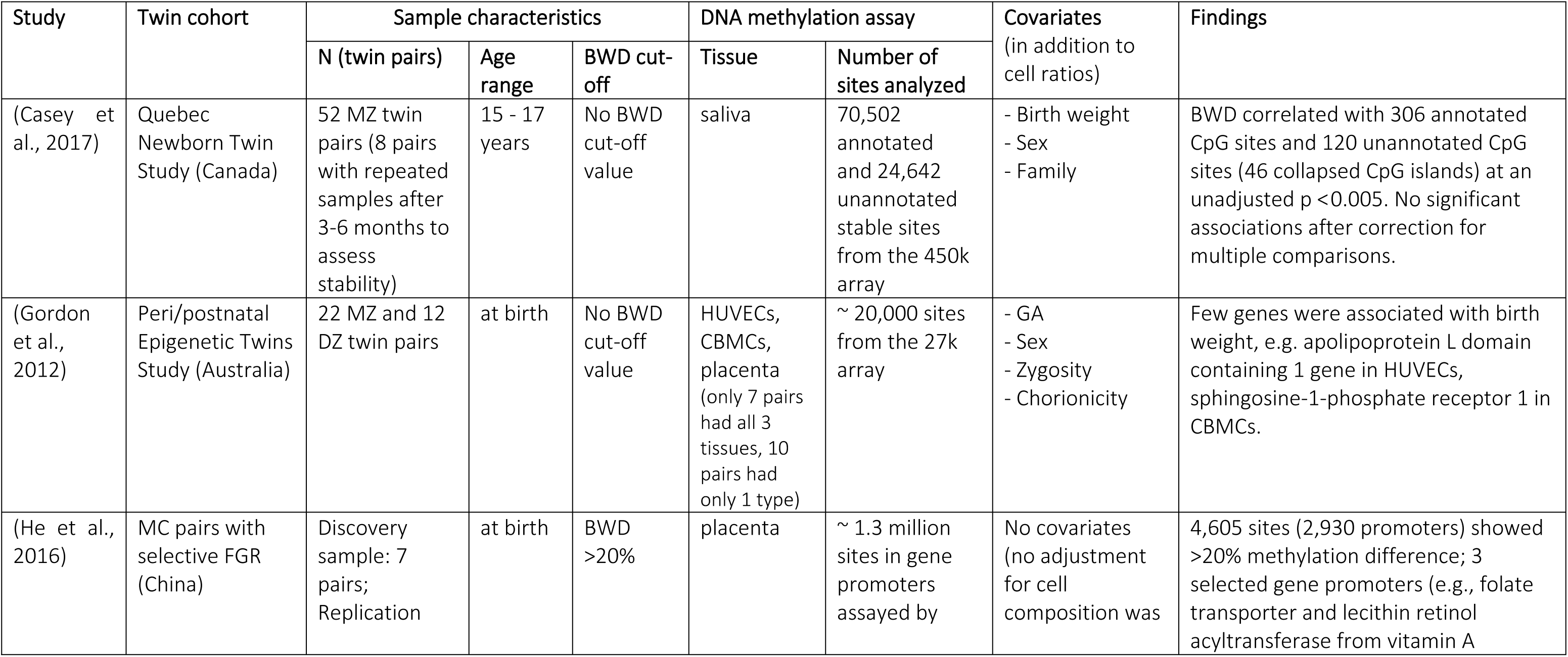
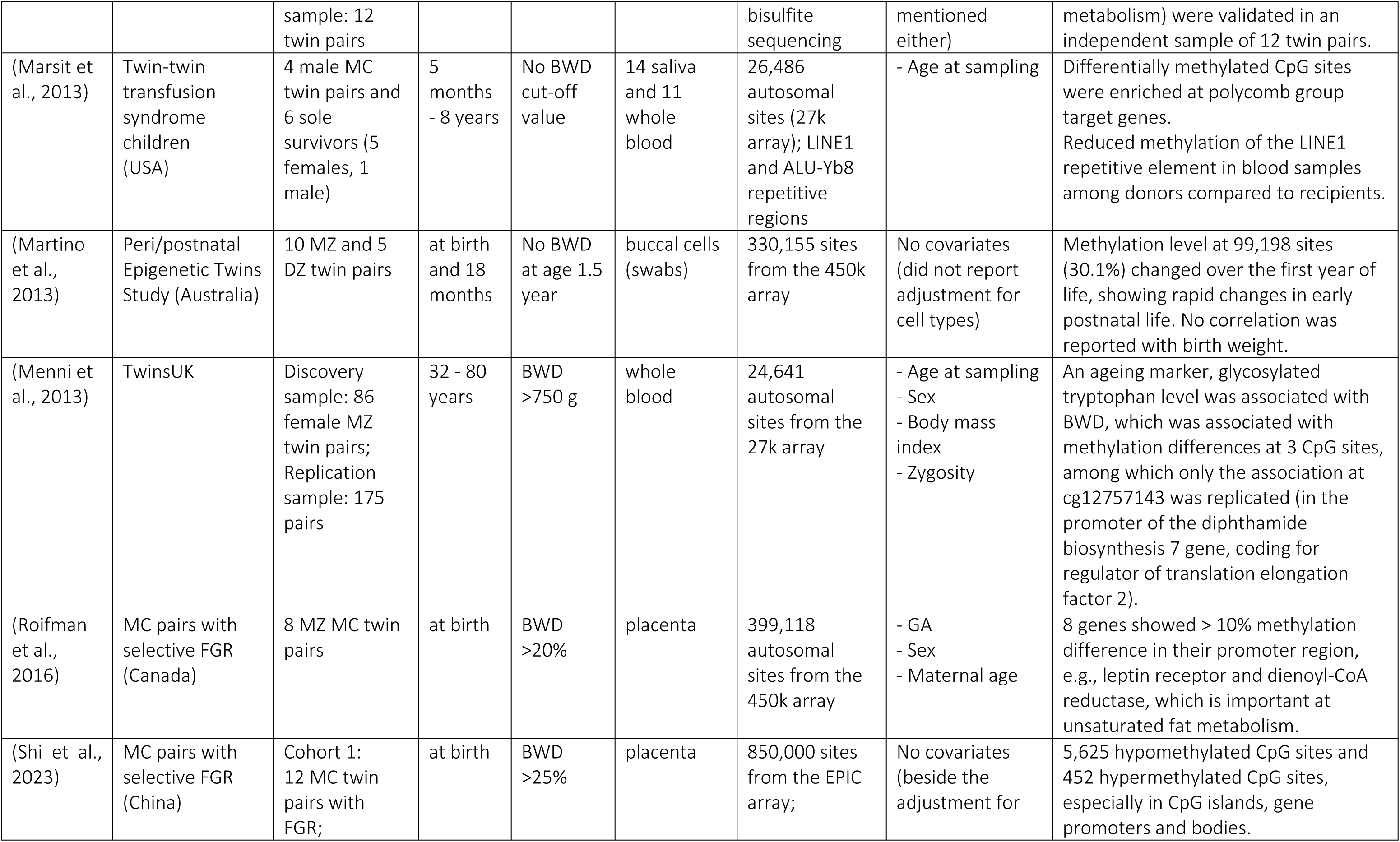
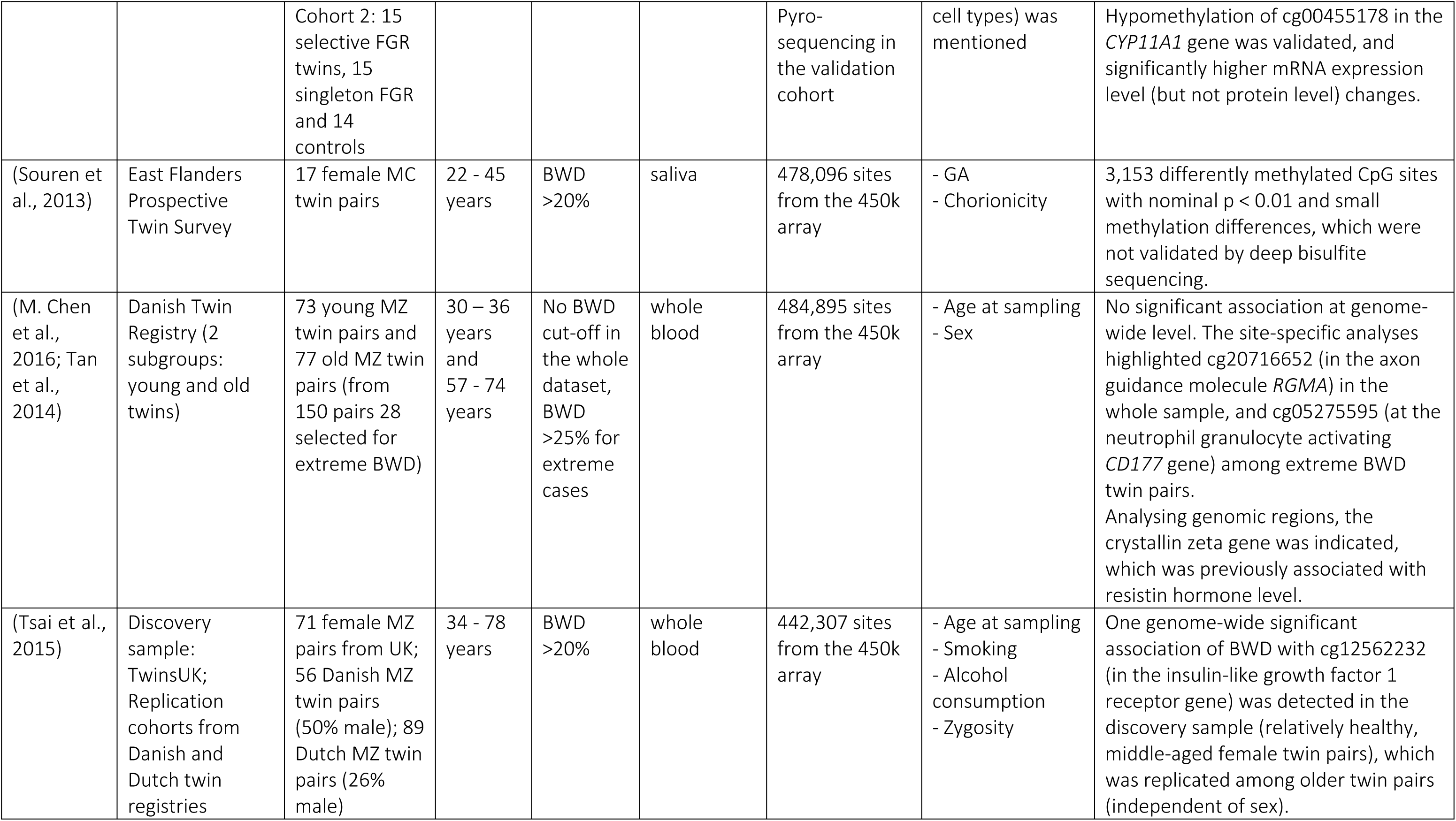
Epigenome-wide twin studies analysing DNA methylation levels and birth weight discordance. In most studies the epigenome-wide data was obtained by the Infinium Human Methylation BeadChip array from Illumina (27k or 450k versions) and adjusted for heterogenous cell composition. Please note that a few studies used overlapping twin cohorts (their descriptions are included in the 2^nd^ column). *Abbreviations:* BWD: Birth weight discordance, CBMCs: cord blood mononuclear cells, GA: gestational age, DZ: dizygotic, HUVECs: human umbilical vein endothelial cells, MC: monochorionic, MZ: monozygotic; FGR: fetal growth restriction.

The earliest epigenetic twin studies used tissues at birth and concentrated on FGR or others medical problems. For example, Bamforth et al. (64) analysed X-inactivation patterns in chorion, amnion, and cord samples of 79 female twin pairs, specifically selected for BWD, discordance for congenital anomalies, TTTS, but no correlation was seen with BWD or with any other clinical outcome. The latest studies rely on more sophisticated methods, which allow detection of smaller differences in DNA (hydroxy)methylation levels, either via candidate gene approach or at the epigenome level. The majority of studies selecting *a priori* candidate genes analysed imprinted regions, e.g., the maternally imprinted insulin-like growth factor 2 gene (*IGF2*) and the paternally imprinted *H19* gene (coding for a tumor-suppressor long non-coding RNA), which have a common imprinting center at the 11p15 chromosomal region (65). Once the specific gene is methylated by the imprinting process, it is inactivated on either the maternal or paternal chromosome in a parent-of-origin effect, e.g., the growth regulator *IGF2* gene is expressed only from the paternal chromosome, because it is imprinted, hence inactivated on the maternal chromosome.

### Candidate gene approach

One of the earliest studies analysed the promoter region of the catechol-O-methyltransferase (*COMT*) gene using buccal cells of 5-year-old healthy MZ twin pairs with high (27-40%) BWD (66). The methylation level of the studied CpG sites showed wide variation among the 12 MZ twin pairs, and there was no correlation with birth weight. Later, Schreiner et al. (67) examined 20 MZ twins at age 3-5 years, who were treated for TTTS before the 27^th^ gestational week. They analysed peripheral blood and saliva samples from the twin pairs and unrelated control children and reported high intra-twin-pair correlation in the methylation level of the *COMT* promoter region in both biological sample types. Importantly, they could not detect any correlation between methylation differences and pre- or postnatal growth patterns. We have to note that these early studies did not control for cell heterogeneity, which can cause substantial differences in tissue-specific methylation patterns both in blood-related and oral samples, especially in alternative promoter regions, such as the one regulating the soluble *COMT* isoform, studied by Mill et al. (66) and Schreiner et al. (67). In this aspect, tissue samples with predominantly one cell type are more comparable. Using placental samples (from which blood was carefully removed by washing steps), Gao et al. (47) reported reduced DNA methylation at the *HERVWE1* gene promoter in the lower birth weight twins within the discordant pairs, compared to their larger siblings, whose methylation values were similar to that of singleton babies born at similar gestational age via cesarean section.

Later on, the majority of epigenetic twin studies using the candidate gene approach focused on the imprinted region on chromosome 11p15 containing the *IGF2* and *H19* genes. Various biological samples from individuals of different ages were used. The first such report analysed neonatal and placental samples collected at birth from the Australian Peri/postnatal Epigenetic Twins Study. They examined multiple tissues, such as cord blood (separated to mononuclear cells and granulocytes), HUVECs and buccal cells from 35 DZ and 56 MZ twins, among whom 19 were DC (68). Substantial differences due to tissue specific methylation patterns were observed, but no strong correlation could be detected with BWD (Spearman correlation coefficients ranged from -0.45 to 0.58). Analysing 4-year-old children previously treated for TTTS, Schreiner et al. (69) could observe only one weak association between BWD and slight increase in methylation at the *IGF2* gene region, more pronounced in saliva than in blood (with Spearman correlation coefficient 0.51). However, no difference was found in the serum IGF2 level between donors and recipients either at birth or at age 4. Furthermore, no correlation was observed between IGF2 serum levels and methylation values using the intra-twin differences. After widening the candidate regions and including IGF2-related developmental genes, such as *IGF2BP1-3* (IGF2-binding proteins 1-3), Córdova-Palomera et al. (70) found a positive association between birth weight and DNA methylation level of two CpG sites in the *IGF2BP1* gene in peripheral blood samples of 17 healthy adult MZ twin pairs (among the 248 preselected sites of the IGF2 pathway). Each kilogram increase in birth weight correlated with approximately 8.3% rise in DNA methylation level of the growth-regulator *IGF2BP1* gene, after controlling for sex and gestational age, as well as for age at sampling (which ranged from 22 to 56 years). Since this RNA binding protein stabilizes *IGF2* transcripts, its reduced expression can result in lower IGF2 level (see more details in Table 2). It has to be emphasized that the rest of the studied sites did not show substantial inter-individual variation (less than 5% of methylation differences), hence they were not informative to reflect non-shared environmental effects.

Taken together, candidate gene twin studies resulted mostly in negative findings, either because of the low variability in the methylation level at the targeted sites or the less sensitive methods used, since we should expect to deal with less than 10% of methylation differences in easily available, heterogenous samples, such as whole blood or saliva samples.

### Epigenome-wide association studies

The first EWAS of BWD used samples collected at the Peri/postnatal Epigenetic Twins Study. Gordon et al. (71) compared different tissues obtained at birth from MZ and DZ twin pairs, and found a few significant associations between DNA methylation levels at genes involved in lipid metabolism and birth weight of twins (for details see Table 3). Afterwards, Martino et al. (72) compared DNA methylation patterns of buccal epithelial cells collected at birth and 18 months later with the wide-spread methylation array type from Illumina (450k BeadChip array). It should be noted that these pilot measurements regarded the biological samples (relatively) homogenous and no adjustment for cell composition was mentioned at their analyses. Their main aim was to estimate the genetic and non-shared environmental effects in the framework of Developmental Origins of Health and Disease theory by studying MZ and DZ twins. Afterwards, only MZ twins were examined with sample sizes ranging from 17 to 150 pairs (see details in Table 3).

In their landmark paper using saliva samples from female MC twin pairs, Souren et al. (73) showed that heterogenous cell composition was driving the largest methylation differences in their epigenome-wide dataset obtained with the Illumina 450k array. After controlling for the buccal epithelial cell ratio (and removing the outlier twin pair, who were the only heavy smokers in the cohort), they showed that BWD was associated with more than 3,000 CpG sites, but only 45 of them had methylation differences > 5%, which was regarded as the threshold for technical variation. Importantly, they applied a different technique, the deep bisulfite sequencing to confirm methylation changes at 8 selected CpG sites. Although the methylation values showed good correlation at 5 out of 8 loci between the two methods, the sequencing data produced much less variation (the 5-7% methylation differences between the discordant twins reduced to ≤ 1 % differences), doubting the biological relevance of the studied CpG sites. In order to support their negative findings, DNA methylation levels at repetitive sequences were also measured, resulting in non-significant differences between the heavy and light co-twin groups. Using bigger sample sizes and younger age range, later EWAS could not detect any significant association either between BWD and salivary DNA methylation patterns (74, 14). Similarly, no significant associations were observed between pre-selected epigenetic markers and birth weight in a sample of 1,040 twins using buccal cells collected at age 6-13 (75). Interestingly, using blood samples of adult twins from the Netherlands Twin Register (collected at age 18-79) weak association was detected between birth weight and DNA methylation score (calculated from Illumina array data of 934 CpG sites indicated by the meta-analysis of 18). However, this combined DNA methylation score explained only 0.39% of the variance (whereas polygenic scores explained 1.52% of the variance) in birth weight (75). For reasons of comparisons, we have to note that much higher variance was explained by combined DNA methylation scores at prenatal maternal smoking (16.9%), and body mass index (6.4%) in the same adult dataset.

Similar to the gene expression studies, few research groups selected extreme cases of BWD to find differentially methylated regions in the genome. The first pilot study examined a special sample of children, survivors of TTTS, treated with laser surgery between 16-24 weeks of gestation (76). Only subtle changes (∼ 5% differences) were detected at specific CpG sites from blood and saliva samples between donors and recipients, but they were more likely situated in genes controlled by polycomb group proteins, highlighting developmentally important regions. Subsequent epigenome-wide studies examined placental tissues from twin pregnancies with selective FGR. Using the most common DNA methylation array, a Canadian group detected at least 10% within-pair methylation differences at 8 genes (77) (see Table 3 for study cohort description). Applying a similar twin design but more in-depth analyses, He et al. (78) found 2,930 differentially methylated promoter regions. Selecting four genes related to specific biological functions (e.g., involved in cell migration, RNA binding, vitamin A metabolism, and folate transportation), methylation differences at three target regions were validated in an independent sample of 12 twin pairs. However, DNA methylation differences at these regulatory regions were not correlated with gene expression differences in the placental tissues from 11 pairs. Compared to normally developing co-twins, the placental shares of growth-retarded twins showed lower general DNA methylation and hydroxymethylation levels (measured by mass spectrometry), supporting the epigenetic differences between discordant twins.

Measuring placental samples of MC growth-restricted fetuses compared the their co-twins, Shi et al. (79) found hundreds of CpG sites differentially methylated. Their analysis of affected pathways revealed dysregulation primarily in steroid hormone biosynthesis, metabolism, cell adhesion, and immune response. Importantly, in an integrative analysis of the promoter methylome of FGR placentas, hypomethylation at the *CYP11A1* gene was highlighted when they compared their own dataset with previously published ones (78, 77). The *CYP11A1* gene codes for a cytochrome *P450* monooxygenase involved in the synthesis of cholesterol, steroids, including vitamin D. It catalyzes the conversion of cholesterol to pregnenolone, the first, rate-limiting step in the synthesis of the steroid hormones. Therefore, up-regulation of this enzyme in placenta can potentially disturb progesterone synthesis and can even trigger oxidative stress, leading to growth restriction.

Most of the EWAS used the Illumina 450k methylation array with easily accessible samples, e.g., saliva (74, 73) or whole blood samples (80–83). Only one of them reported a differentially methylated CpG site at the genome-wide significance level using a female MZ twin sample with BWD >20% selected from the TwinsUK cohort. Interestingly, this site in the insulin-like growth factor 1 receptor (*IGF1R*) gene could be detected only with quantitative analysis using BWD as a continuous variable (83). Using the same TwinsUK sample in their ageing-related metabolic analyses, Menni et al. (80) detected significant associations between an ageing marker, the glycosylated tryptophan level and BWD at 3 genes (diphthamide biosynthesis 7, endothelin 2, galactosidase beta 1 like 3), among which one was replicated in their combined dataset of 522 individuals. Association between glycosylated tryptophan metabolite and DNA methylation level at cg12757143 in the promoter region of the diphthamide biosynthesis 7 gene pointed to the involvement elongation factor 2 regulation, which is important not only in protein synthesis but also in the regulation of cell proliferation and migration. Importantly, similar associations could be detected in other, independent twin samples (see Table 3 for details).

Using a relatively big (and not preselected) twin cohort, Tan et al. (82) conducted EWAS on 150 MZ pairs using both quantitative analyses (with BWD as a continuous variable) and qualitative analyses (comparing the larger and smaller co-twins), but could not detect any genome-wide significant associations. In the same dataset of the Danish twin cohort, M. Chen et al. (84) searched for differentially methylated gene regions. One BWD-associated chromosomal region (1p31) could be detected in their quantitative analyses, involving a metabolism-related gene, the crystallin zeta gene coding for a NADPH-dependent quinone reductase, which was previously associated with resistin hormone levels in a genome-wide association study (85), and subsequently linked with insulin resistance (an important metabolic marker of prediabetes, showing the reduced sensitivity to insulin in the fat and muscle tissues).

In order to reveal biologically relevant genes or pathways affected by BWD, we checked the gene names provided by the different groups listing top CpG sites associated with either birth weight as a continuous variable in singleton cohorts or with BWD by comparing heavy and light co-twins. Only a few overlapping genes were detected, such as the PPARG coactivator 1 beta (*PPARGC1B*) involved in fat oxidation and regulation of energy expenditure, or the RAS p21 protein activator 3 (*RASA3*) participating in one of the important pathways regulating cell growth and division. However, many epigenetic twin studies highlighted genes involved in steroid synthesis, glucose metabolism, and lipid oxidation.

## Discussion

This review aimed to summarize the state of research about epigenetic modifications associated with BWD in twin pregnancies. We have to mention that birth weight is predominantly driven by shared environmental factors, mostly by gestational age and maternal factors. After controlling for gestational age, genetic and unique environmental effects accounted for roughly equal shares of the variation in birth weight (20-25%), although there were some cultural-geographical differences (86). Therefore, researchers tried to explore these unique environmental effects and their potential underlying biological mechanisms. As in many complex, polygenic traits, birth weight is controlled by hundreds of genetic variants and possibly similarly high numbers of epigenetic modifications, each contributing a small percentage to the variance. Confirming the relevance and importance of such variants, meta-analyses with thousands of study participants are needed (see for example 18).

Using the candidate gene approach, researchers have focused on genes with profound roles in fetal growth, oxygen-regulation, glucose metabolism and neurodevelopment. The genome-wide expression analysis by Gordon et al. (61) showed significant associations between measures of BWD and metabolism-related genes, like Leptin Receptor Overlapping Transcript (*LEPROT*), adrenoreceptor beta 3, and glucose transporter 3. Candidate gene analyses also supported the involvement of leptin signaling pathway and glucose transport. It should be noted that most expression studies used less than 30 samples per group, therefore, they can be regarded as small-scale pilot studies. Much bigger sample sizes were available at the DNA methylation studies, but the results obtained from recent EWAS are still inconsistent. It might be because of the weaker functional relevance, since DNA methylation changes by themselves do not necessarily reflect gene expression changes, as it has been demonstrated at the imprinted *H19* or *IGF2* genes (87). Importantly, most studies could not detect any significant differences in DNA methylation patterns between the heavy and light co-twins using categorical analyses. The most promising findings were obtained from blood samples of MZ twins with BWD > 20% using quantitative analysis (84, 83). Epigenetic profiles of saliva samples showed much more variability (73).

Reasons for discrepancies may lie in the study design. First, the age of sampling, i.e., the time period at which methylation measurements were taken, differs from study to another. For example, DNA methylation was examined after delivery (71), during adolescence (74) or in adulthood (84, 73, 81, 83). The results are therefore not always comparable, since DNA methylation is not fixed at many CpG sites over the life span, although there are certain sites with stable methylation pattern, as shown by longitudinal studies with multiple timepoints (e.g., 88, 20). Second, DNA methylation level was measured mostly only once, and from only one tissue type, using Illumina arrays which have preselected CpG sites of the human genome. Third, potential differences may also stem from the type of tissues used in these studies (e.g., saliva, blood, neonatal tissues) since cellular heterogeneity was not controlled in the earlier studies and each cell type has its unique epigenetic landscape that likely reflects its specific function and response to environmental exposures. It should be also noted, that most of the EWAS included all ∼ 480 thousand CpG sites from the 450k methylation array, although the majority of CpG sites on the Illumina arrays are not variable between individuals, which potentially had a major effect at the statistical analyses (therefore the analysed numbers of CpG sites are indicated Table 3). Newer types of analyses on merged, large-scale datasets are needed for convincing results.

Epigenetic regulation of growth-related genes in the offspring may be a relevant mechanism through which adverse prenatal environment is associated with low birth weight. The literature also shows similar trends in studies of singletons. In fact, most researchers that examined singletons showed that birth weight (adjusted or not for gestational age) was associated with DNA methylation patterns at birth or later in life at certain CpG sites involved in growth or in embryonic development (18). Newer studies did not replicate the previously highlighted genes, however, they used much smaller sample sizes, hence they were underpowered to detect few percentages of epigenetic difference between low and high birth weight infants (89, 90). Taken together, this review showed how DNA methylation patterns associated with BWD are characterized by variability in methodology (e.g. sampling time, tissue type and DNA methylation measure used), likely contributing to the inconsistency of results reported.

We can potentially expect a couple of reliable findings on the genome-wide significance level in the near future, as there are many large twin studies with EWAS data using peripheral blood samples collected from young adults (age 18-36 years), such as the UK-based E-Risk Twin Study (N = 1629), the Finnish Twin Cohorts (N = 757), and Netherlands Twin Register (N = 2059) (see the discovery and replication samples of a recent meta-analysis by 91). These twin samples are part of the project “Unraveling gene-environment interplay to inform Treatment and Intervention strategies” which is now in the analytical phase. Newer, more sophisticated analyses could detect important epigenetic alterations from early developmental stages, which can stay stable in our genome and can serve as useful epigenetic markers later in life (at the potential disease prognostic stage). Although it is easier to collect thousands of adult participants with retrospective data, these analyses should control for larger environmental effects first, such as current metabolic and behavioral variables (e.g., body mass index, smoking). Prospective studies with reliable, detailed information about perinatal factors are needed to highlight individual DNA methylation sites with small effects.

Finally, we would like to note some gaps in the literature, which can be addressed by future studies. For example, researchers could examine several attractive candidate genes with known involvement in fetal growth (e.g., *IGF2*, *LEPROT, ADRB3*, *GLUT3*) in two or more tissues and developmental stages (e.g., in placenta, cord blood or umbilical vein cells, and in child and adult blood or saliva samples), which can help finding stable epigenetic modifications associated with DNA methylation differences between the co-twins. Further, most studies conducted to date have been cross-sectional in design and relied on single epigenetic measures using samples collected only once at different time points, like delivery, early childhood, adolescence or adulthood. Longitudinal studies with several data collection points for epigenetic measures would be useful to comprehensively and prospectively investigate the long-term effects of prenatal environmental exposures indicated by BWD. In addition, few of the association studies used extreme BWD pairs of twins (i.e. BWD > 30-40%). Our understanding may be improved by including twins with wide range of BWD. Lastly, looking at potential confounding factors, many studies did not control for some major confounders, such as chorionicity, gestational age, zygosity and sex, which can likely account for a large proportion of the observed association between prenatal environmental exposures, BWD, and epigenetic changes as suggested by researchers (71, 73).

## Conclusion

This review found several studies which showed that prenatal environment experienced *in utero* by twins can be associated with BWD through altered gene expression, particularly at genes involved in pathways affecting fetal growth, which in turn can affect birth weight. However, these studies had small sample sizes and did not include replication cohorts. Larger size twin studies are available examining BWD and DNA methylation patterns, but have not provided consistent evidence yet, possibly due to the small methylation differences in tissue samples with heterogenous cell compositions. In order to reveal true associations, future epigenome-wide analyses should pay attention to several confounding parameters to be controlled for (e.g., smoking), and include sample size with sufficient power to detect a few percentages of methylation differences.

## Acknowledgement

This project was supported by a grant from the Swiss Government Excellence Scholarships (2018.0801, 2018) and the Swiss National Science Foundation (P1FRP1_199872, 2021) awarded to DLW. We are grateful to Linda Booij for helping and supporting this review. The work was facilitated by the Canadian Institutes of Health Research (CIHR) grant # 162306.

